# Machine Learning Based Identification and Characterization of Mast cells in Eosinophilic Esophagitis

**DOI:** 10.1101/2023.10.25.563471

**Authors:** Simin Zhang, Julie M. Caldwell, Mark Rochman, Margaret H. Collins, Marc E. Rothenberg

## Abstract

**Background:** Eosinophilic esophagitis (EoE) is diagnosed and monitored using esophageal eosinophil levels; however, EoE also exhibits a marked, understudied esophageal mastocytosis.

**Objective:** Using machine learning, we localized and characterized esophageal mast cells to decipher their potential role in disease pathology.

**Methods:** Esophageal biopsy samples (EoE, control) were stained for mast cells by anti-tryptase and imaged using immunofluorescence; high-resolution whole tissue images were digitally assembled. Machine learning software was trained to identify, enumerate, and characterize mast cells, designated Mast Cell-Artificial Intelligence (MC-AI).

**Results:** MC-AI enumerated cell counts with high accuracy. During active EoE, epithelial mast cells increased and lamina propria (LP) mast cells decreased. In controls and EoE remission patients, papillae had the highest mast cell density and negatively correlated with epithelial mast cell density. Mast cell density in the epithelium and papillae correlated with the degree of epithelial eosinophilic inflammation, basal zone hyperplasia, and LP fibrosis. MC-AI detected greater mast cell degranulation in the epithelium, papillae, and LP in EoE patients compared with control individuals. Mast cells were localized further from the basement membrane in active EoE than EoE remission and controls individuals but were closer than eosinophils to the basement membrane in active EoE.

**Conclusion:** Using MC-AI, we identified a distinct population of homeostatic esophageal papillae mast cells; during active EoE, this population decreases, undergoes degranulation, negatively correlates with epithelial mast cell levels, and significantly correlates with distinct histologic features. Overall, MC-AI provides a means to understand the potential involvement of mast cells in EoE and other disorders.

**Clinical Implication:** We have developed a methodology for identifying, enumerating, and characterizing mast cells using artificial intelligence; this has been applied to decipher eosinophilic esophagitis and provides a platform approach for other diseases.

**Capsule Summary:** A machine learning protocol for identifying mast cells, designated Mast Cell–Artificial Intelligence, readily identified spatially distinct and dynamic populations of mast cells in EoE, providing a platform to better understand this cell type in EoE and other diseases.

## Introduction

Eosinophilic esophagitis (EoE) is a food-induced allergic inflammatory disease of the esophagus. Although rare, the prevalence of EoE is increasing with current estimates being 56 cases per 100,000 persons^1,2^. Active disease is determined by upper endoscopy, and the pathologic diagnosis is defined by at least 15 eosinophils per high-power field (eosinophils/hpf [eos/hpf]) identified on esophageal biopsies, yet recent data indicate that eosinophils are not involved in the clinical manifestations of eosinophilic gastrointestinal diseases, which calls attention to the potential pathogenic role of other cell populations^3–5^. Notably, there is concurrent mastocytosis and some evidence of mast cell degranulation in the esophageal epithelium^6–8^. Furthermore, the density of mast cells correlates more highly with certain symptoms and histologic features than eosinophils, suggesting that mast cells are clinically relevant and involved in disease pathogenesis, including esophageal epithelial barrier disruption^9–13^. Unlike eosinophils, which do not reside in the esophageal epithelium in healthy states, mast cells exist in small numbers in the esophageal epithelium and in the esophageal lamina propria (LP) during homeostasis^8^. Furthermore, in active disease, they produce inflammatory mediators such as IL-13 which has been shown to be an essential driver of EoE ^14,15^.

The need for specialized staining, such as anti-tryptase, to visualize mast cells and the time-intensive nature of manual counting hinder their assessment in the esophagus. Additionally, morphometric approximation of mast cells is difficult due to their pleomorphic nature. Herein, we developed a machine learning segmentation protocol for localizing and characterizing esophageal mast cells, including their degranulation, and we applied it to further understand the potential role of mast cells in EoE.

## Methods

### Patients

The Cincinnati Children’s Hospital Medical Center (CCHMC) institutional review board approved all human studies, and subjects provided written consent. Active EoE was defined as patients whose esophageal biopsy had peak eosinophil count ≥15 eos/hpf. EoE remission was defined as patients with a history of EoE diagnosis whose current esophageal biopsy had a peak eosinophil count <15 eos/hpf. Control patients were defined as those who underwent esophageal biopsy with no prior history of EoE and had a peak eosinophil count ≤1 eos/hpf.

### Immunofluorescence (IF)

Human esophageal biopsies from the Cincinnati Center for Eosinophilic Disorders (CCED) biorepository were randomly selected without consideration of sex, race, atopic status or treatment. Formalin-fixed, paraffin-embedded slides were first deparaffinized with xylene, subjected to graded ethanol washes and subsequent antigen retrieval with antigen retrieval buffer under pressurized conditions, washed in PBS & Triton (0.2%), and blocked in 10% donkey serum in PBS. Biopsies were incubated with primary antibodies (1:50) (anti-tryptase, mouse IgG1 monoclonal antibody (Biolegend)) overnight at 4°C; isotype antibodies (anti-IgG1) were used as negative controls. Biopsies were then incubated for 1 hour with Hoechst for nuclear staining and fluorescent secondary antibodies (donkey anti-mouse Alexa Fluor 488). Sequential, partially overlapping images were taken on a Nikon A1R LUN-V inverted confocal microscope at 40X magnification and stitched together into multipoint, whole biopsy images.

### Mast cell training and validation using Segment.ai

NIS.ai is a machine learning add-on package for Nikon Elements (Nikon, Tokyo, Japan) that allows automation of processes that were previously difficult to achieve by conventional morphometric methods. One feature of NIS.ai is that Segment.ai uses a pre-developed machine learning algorithm that can be trained to recognize structures of interest. Segment.ai software was trained to recognize mast cells according to guidance from the manufacturer. Mast cells were identified by the presence of any tryptase staining around an identifiable nucleus. For Segment.ai to optimally identify mast cells, high-resolution confocal images were taken at 40X hpf, and then Elements software was used to digitally stitch these images together to assemble a whole biopsy section composite image on which Segment.ai was then applied. To train Segment.ai to identify mast cells, mast cells were hand traced (S.Z.) in whole biopsy images (from 1 control patient and 1 patient with active EoE) and used for the training set. Subsequently, 17 biopsies (n = 8 controls; n = 9 active EoE) were employed as a test set to compare manual counting– and Segment.ai-based mast cells counts. To capture mast cells more accurately, post Segment.ai–identified mast cells were modified with “close” and “dilate” functions, and the area of mast cells was restricted to 10–200-µm sizes.

### Mast cell density and degranulation

Area measurements were first calculated with a general analysis formula based on nuclear staining using Elements software (Nikon, Tokyo, Japan). Low-level whole tissue autofluorescence at 651 nm wavelength was used to manually separate the epithelium, papillae, and LP. Esophageal papillae were defined as any LP that projected past two points of basement membrane (bisectional) and areas of LP that were surrounded by epithelium (cross-sectional). To determine spatial distribution, the distance from the center of MC-AI–identified mast cells to the manually identified basement membrane was measured.

Subsequently, a separate protocol was trained on 1 active EoE biopsy section to detect degranulated mast cells (hand traced by S.Z.). Degranulated mast cells were identified as mast cells with tryptase that were not well circumscribed. This was termed degranulated MC-AI. A second method used for identifying degranulation was using fluorescence intensity thresholding to identify all tryptase staining on biopsy sections and then subtracting mast cells identified by the original MC-AI protocol to approximate extracellular tryptase. A third method used the mean fluorescence intensity (MFI) of MC-AI–identified mast cells to approximate mast cell degranulation, as degranulation of mast cells should result in decreased tryptase MFI.

### Eosinophil density

Eosinophil autofluorescence in IF at 544 nm wavelength was observed if the channel was left open without any additional antibody staining. This autofluorescence was used to identify eosinophils for enumeration. Eosinophils identified in one active EoE biopsy were hand traced (S.Z.), and Segment.ai was trained to recognize these eosinophils.

### Histology Scoring System (HSS)

The HSS evaluates 8 histologic features of eosinophilic esophagitis, specifically eosinophilic inflammation (EI), basal zone hyperplasia (BZH), eosinophilic abscesses (EA), eosinophilic surface layering (ESL), dilated intercellular spaces (DIS), surface epithelial alteration (SEA), dyskeratotic epithelial cells (DEC), and LP fibrosis (LPF)^16^. The HSS contains a 4-point scale from 0 (normal) to 3 (most severe/extensive) for grade (severity) and stage (extent) of each feature, which totals a maximum score of 24 for grade or stage. The sum of the scores is divided by 24 for the final grade or stage score if all features were evaluable. If all features were not evaluable, the observed sum is divided by the number of evaluable features and then multiplied by 3.

### Statistical analyses

GraphPad Prism 9.5 (GraphPad Software, San Diego, California) was used for all statistical analyses. Linear regression was used for correlation analyses. The Mann-Whitney U test was used for non-parametric comparisons between 2 categories.

## Results

### Machine learning identification of mast cells reveals dynamic mast cell density as a function of location and EoE disease activity

Human esophageal biopsy sections were stained with a nuclear dye (Hoechst), and anti-tryptase antibody and digital images were stitched together to identify mast cells manually and by Segment.ai (Figure 1A). MC-AI–identified mast cells positively correlated with manually identified mast cells (n = 17 biopsy sections, R^2^ = 0.95, p<0.0001) (Figure 1B).

**Figure 1.**
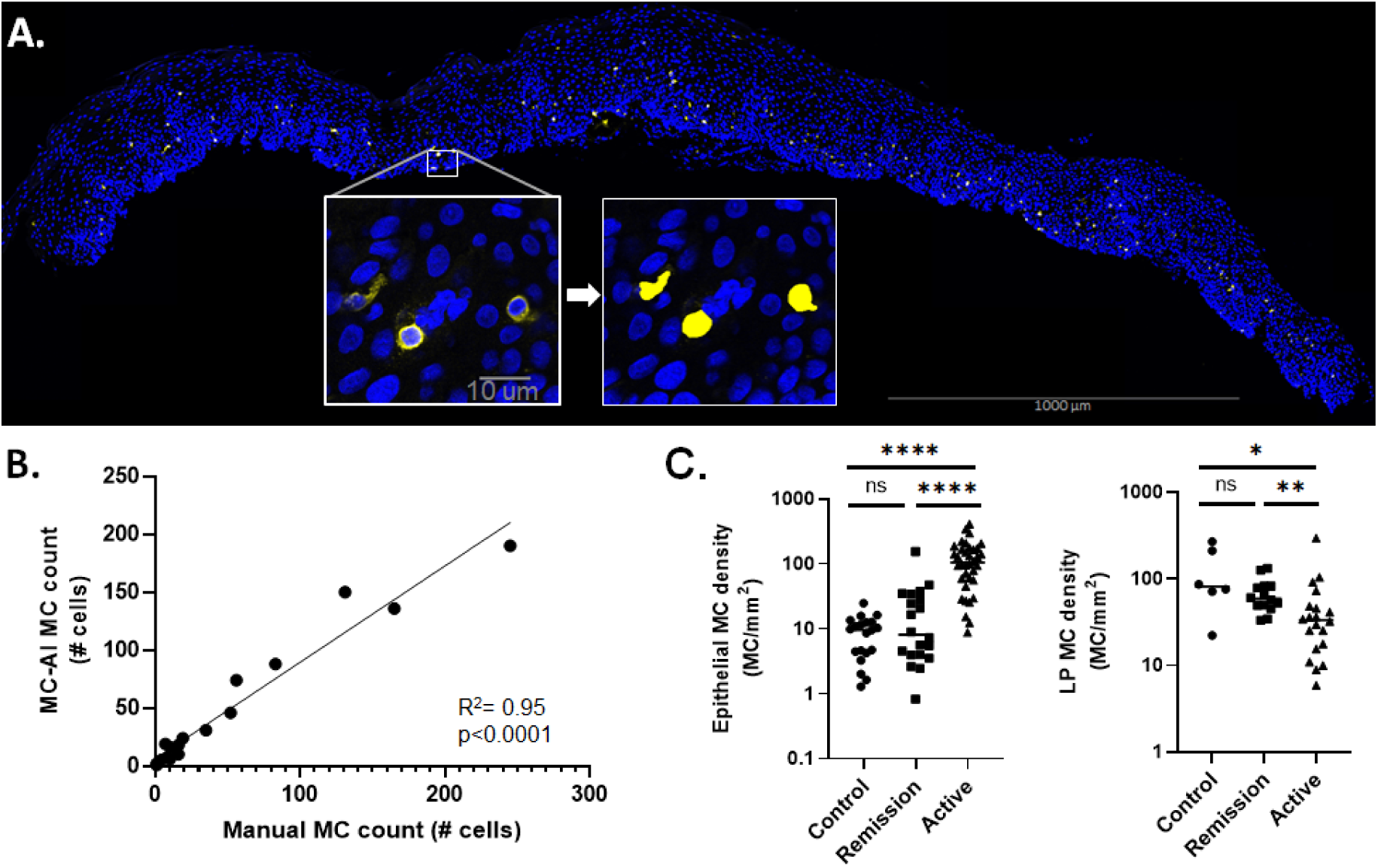
MC-AI method and MC-AI–identified MC density as a function of disease state. A. Digitally stitched whole biopsy immunofluorescence of human esophageal tissue, multiple 40X high-powered fields (hpf). Nuclear (blue) and tryptase (yellow) staining for mast cells (MCs). Image from one representative biopsy section; 40X hpf showing MCs stained for tryptase (left) and Segment.ai (MC-AI)–identified MCs (right). B. Correlation between manual and MC-AI counting of epithelial MCs (simple linear regression), n = 17 biopsy sections. C. Epithelial MC density in control (n = 21 patients), EoE remission (Remission, n = 20 patients), and active EoE (Active, n = 36 patients) (left) and lamina propria (LP) MC density in control (n = 6 patients), EoE remission (n = 14 patients), and active EoE (n = 20 patients) (right); markers represent individual patients, and bars represent the mean. Statistical differences between groups *p < 0.0332, **p < 0.0021, ***p < 0.0002, ****p < 0.0001, ns = not significant.

MC-AI enabled quantification of mast cell density as a function of EoE disease activity (Table 1). There was an increase in epithelial mast cell density in active EoE compared to control (p < 0.0001) and EoE remission (p < 0.0001) (Figure 1C). There was a decrease in LP mast cell density in active EoE compared to control (p = 0.04) and EoE remission (p = 0.004).

**Table 1.** Mast cell density (cells/mm^2^) by location and disease state.

|  | **Control** | **Remission** | **Active** | **p value** |
| --- | --- | --- | --- | --- |
| **Epithelium** | 9 (mean) | 22 | 124 | p < 0.0001 |
|  | 10 (median) | 8 | 104 |  |
|  | 0-25 (range) | 1-155 | 9-409 |  |
| **n** | 21 | 20 | 36 |  |
| **Papillae** | 536 | 291 | 125 | p < 0.0001 |
|  | 591 | 301 | 79 |  |
|  | 0-1,386 | 0-601 | 0-737 |  |
| **n** | 19 | 18 | 35 |  |
| **LP** | 123 | 68 | 49 | p = 0.0069 |
|  | 81 | 59 | 33 |  |
|  | 22-269 | 33-132 | 6-294 |  |
| **n** | 6 | 14 | 20 |  |

Active, active eosinophilic esophagitis; Control, non-eosinophilic esophagitis; LP, lamina propria; n, number of biopsy sections; Remission, eosinophilic esophagitis in remission.
Kruskal-Wallis test used for calculation of p value.

### Papillae-resident mast cells and their densities as a function of EoE disease activity

Human esophageal tissue was found to have low-level autofluorescence at 651 nm wavelength that distinguished the epithelium from the LP (S. Figure 1). The LP of the esophagus had papillae, defined as projections of LP into the epithelium. E-cadherin staining, an epithelial marker, showed staining only in the epithelium, which confirmed the separation of the epithelium from the papillae and LP as seen with autofluorescence (Figure 2A). The low-level autofluorescence facilitated tracing of esophageal papillae, allowing separation of mast cells residing in the epithelium from the papillae (Figure 2B). The compartment with the highest mast cell density in control and EoE remission biopsy sections was the papillae, followed by the LP and then the epithelium (Figure 2C). In active EoE, there was no difference in mast cell density between the epithelial and papillary compartments (p = 0.34). There was a decrease in the papillary mast cell density in control versus EoE remission (p = 0.02) and in control versus active EoE (p < 0.0001). There was a negative correlation between epithelial and papillary mast cell density (R^2^ = 0.4, p < 0.0001) (Figure 2D).

**Figure 2.**
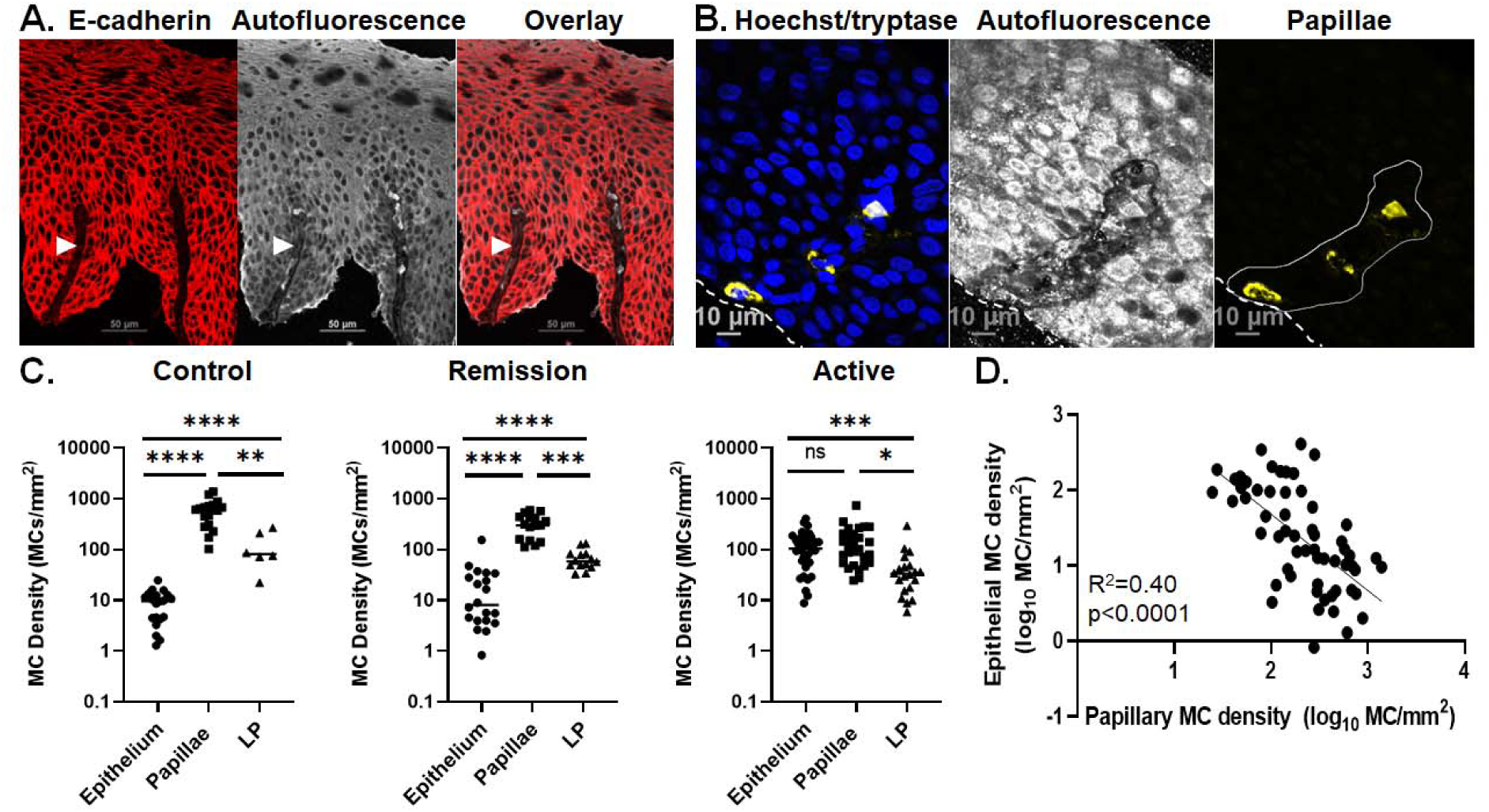
Papillae and dynamic changes of MC-AI–identified papillary MC density as a function of disease state. A. Immunofluorescence of E-cadherin (left, red) representing esophageal epithelium, low-level autofluorescence at 640-nm wavelength (middle, white), and their overlay (right). Arrowhead: papillae. B. Immunofluorescence of esophageal epithelium with nuclear (blue) and mast cell (MC) tryptase (yellow) staining (left), low-level autofluorescence at 640 nm (middle, white), and manual tracing of papillae overlaid with MC tryptase (yellow) (right). A-B images are crops of digitally stitched images of composite overlapping 40X images of one representative biopsy section each. C. MC density in control papillae (n = 19 biopsy sections), epithelium (n = 21), and lamina propria (n = 6) (left); EoE remission (Remission) papillae (n = 18), epithelium (n = 20), and lamina propria (n = 14) (middle); and active EoE (Active) papillae (n = 35), epithelium (n = 36), and lamina propria (n = 20) (right). Markers represent individual biopsy sections, and bars represent the mean. D. Correlation of log-transformed values between papillary and epithelial MC density; markers represent individual biopsy sections. Statistical difference between groups; *p < 0.0332, **p < 0.0021; ***p < 0.0002, ****p < 0.0001, ns = not significant.

We aimed to determine whether papillary mast cells are potentially mast cell progenitors, perhaps derived from the circulation, as papillae are highly vascular structures^17^. We hypothesized that if papillary mast cells were progenitors entering from the vasculature present in the papillae, then some of the papillary mast cells would be CD34 positive. Human esophageal biopsy sections were co-stained with anti-tryptase and anti-CD34, a marker for hematopoietic progenitors, including mast cell progenitors^18,19^ (Figure S2A). Out of the 4 biopsy sections, 9 papillary mast cells were identified, none of which were CD34 positive (thus, at least ∼90% of papillary mast cells do not express CD34 (Figure S2B). Epithelial mast cells (n=44) were also CD34 negative (data not shown).

### Mast cell density associates with EoE histologic features

Subsequently, we aimed to determine whether mast cell density in the different compartments (epithelium papillae, LP) was associated with disease pathology. To test this hypothesis, epithelial, papillary, and LP mast cell density were correlated with HSS grades (Figure 3A) and stages (Figure 3B) in patients across all disease states (active EoE, EoE remission, control). Across all disease states, epithelial mast cell density was found to be associated with HSS features, including eosinophilic inflammation grade (p < 0.0001) and stage (p < 0.0001), basal zone hyperplasia grade (p < 0.0001) and stage (p < 0.0001), eosinophilic abscess stage (p = 0.029), LP fibrosis grade (p = 0.0076) and stage (p = 0.011), and total HSS score grade (p < 0.0001) and stage (p < 0.0001). Across all disease states, papillary mast cell density was found to be negatively associated with HSS features, including eosinophilic inflammation grade (p = 0.0003) and stage (p = 0.0009), basal zone hyperplasia grade (p = 0.0042) and stage (p = 0.032), LP fibrosis grade (p = 0.031) and stage (p = 0.022), and total HSS score grade (p = 0.0077) and stage (p = 0.0029). In contrast, LP mast cell density did not significantly correlate with any HSS features. Of note, eosinophil density correlated with eosinophilic inflammation grade (p = 0.0056) and stage (p = 0.0003), basal zone hyperplasia grade (p = 0.0016) and stage (p = 0.0041), total HSS score grade (p = 0.018) and stage (p = 0.0071); papillae eosinophil density correlated with eosinophil surface layering grade (p = 0.012) and stage (p = 0.0001); lamina propria eosinophil density correlated with basal zone hyperplasia grade (p = 0.029) and eosinophil surface layering grade (p = 0.002) and stage (p = 0.002) (Figure S3A).

**Figure 3.**
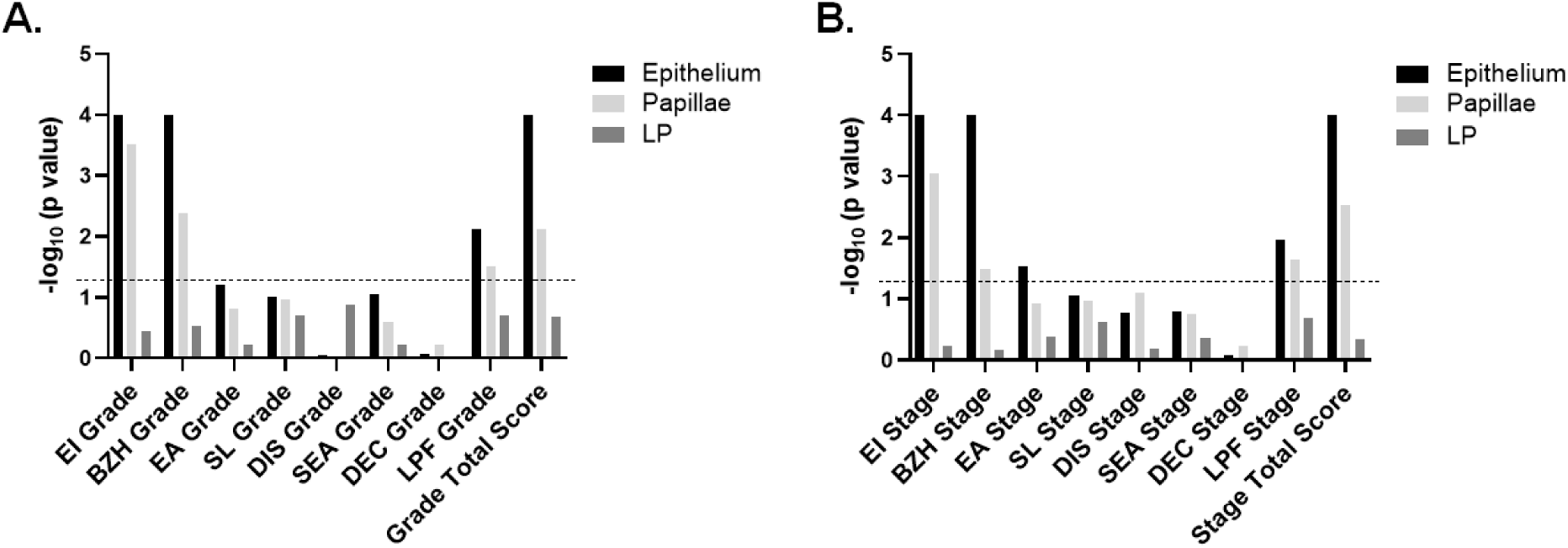
Correlation of MC-AI–identified MC density and Histology Scoring System (HSS) scores across all disease states (-log 10–adjusted p values). A. Histology Scoring System (HSS) grade scores. B. HSS stage. Dotted line represents -log 10–adjusted p value of 0.05. Simple linear regression was used for analysis. LP, lamina propria, EI, eosinophil inflammation, BZH, basal zone hyperplasia, EA, eosinophil abscess, SL, eosinophil surface layering, DIS, dilated intercellular spaces, SEA, surface epithelial alteration, DEC, dyskeratotic epithelial cells, LPF, lamina propria fibrosis.

### Evaluating mast cell degranulation with MC-AI and extracellular tryptase

We next investigated mast cell degranulation in esophageal tissue, aiming to initially train Segment.ai to recognize degranulated mast cells. We manually outlined degranulated mast cells in a single biopsy section, trained Segment.ai, and designated this protocol Degranulated MC-AI (Figure 4A). Degranulated mast cell counts for the trained degranulated MC-AI protocol correlated with manual counts (R^2^ = 0.71, p = 0.0003, n = 13), although the counts were higher in the Degranulated MC-AI protocol (Figure 4B). Degranulated mast cell density, as identified by Degranulated MC-AI, was higher in EoE than control (Figure 4C, p = 0.002, n = 6 EoE, 7 control).

**Figure 4.**
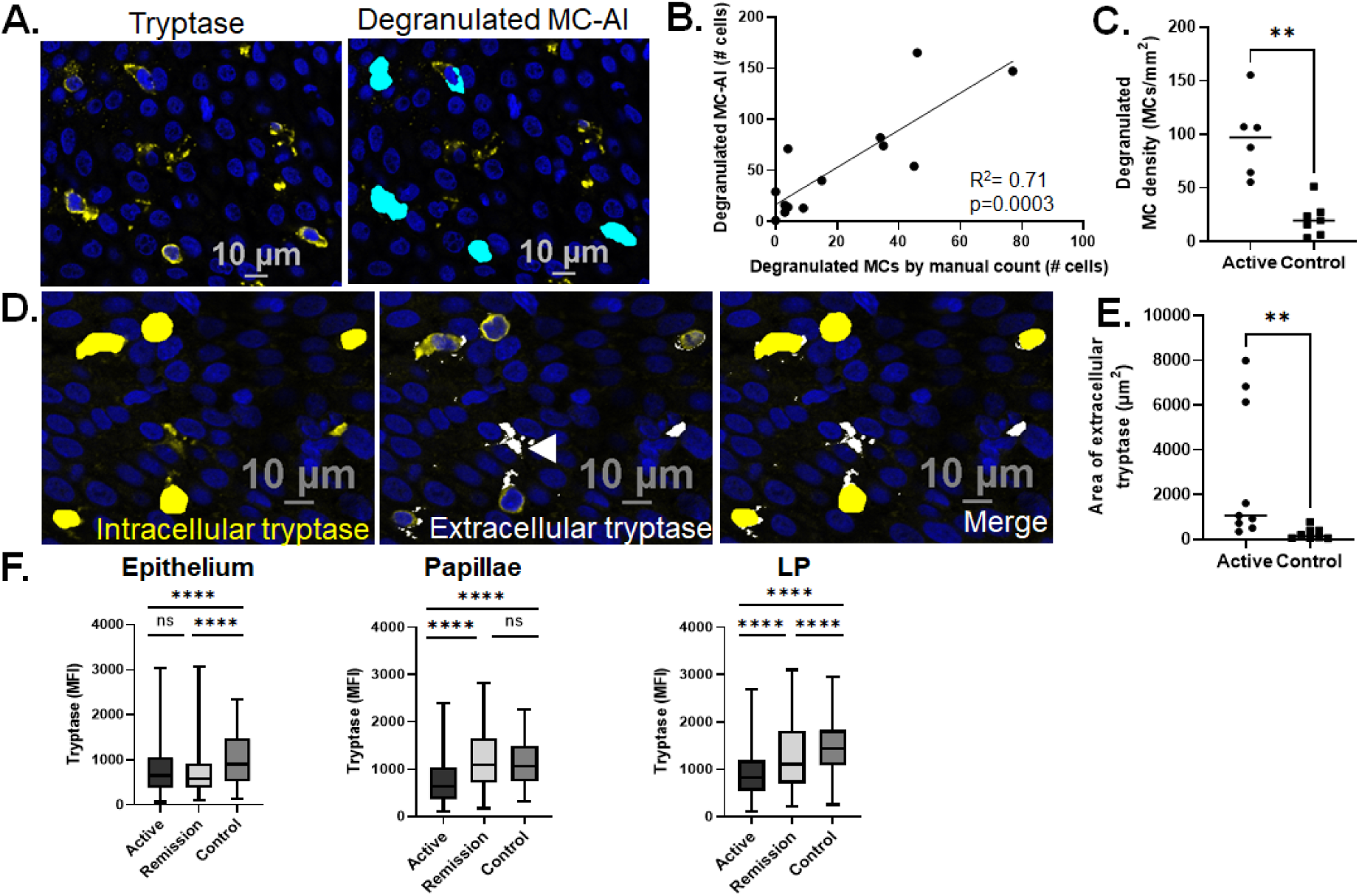
Degranulated MC-AI. A. Crop of digitally stitched whole biopsy immunofluorescence of human esophageal tissue, multiple 40X high-powered fields (left). Nuclear (blue) and tryptase (yellow) staining for mast cells (MCs). Degranulated MC-AI– identified degranulated MCs (cyan) (right). B. Correlation between manual and Degranulated MC-AI–identified degranulated MCs (simple linear regression, n = 13 biopsy sections). C. Degranulated MC-AI–identified degranulated MC density in active EoE (Active, n = 6 biopsy sections) and control (n = 7). D. MC-AI–identified MCs were used to identify intracellular tryptase (yellow, left); tryptase was identified by fluorescence intensity thresholding of anti-tryptase (right), and intracellular tryptase (left) was subtracted to identify extracellular tryptase (white; arrowhead; middle); merged areas of intracellular and extracellular tryptase (right). E. Area of extracellular tryptase per biopsy section as identified using method in D in active EoE (n = 9 biopsy sections) and controls (n = 8). C and E. Markers represent individual patient biopsy sections; bars represent the median. F. Tryptase mean fluorescence intensity (MFI) of individual MCs in esophageal epithelium in active EoE (n = 2,858 MCs), EoE remission (Remission, n= 445), and control (n= 119) patients (left); in papillae in active EoE (n = 95), EoE remission (n = 69), and control (n = 61) patients; and in lamina propria (LP) in active EoE (n = 202), EoE remission (n = 325), and control (n = 174) patients. Box represents 25-75% of values, lines represent 1-100%. Statistical differences between groups *p < 0.0332, **p < 0.0021, ***p < 0.0002, ****p < 0.0001, ns = not significant.

An alternative method was also used to capture degranulation. Intensity thresholding was used to identify all tryptase staining on biopsy sections; the mast cell areas as identified by MC-AI were subtracted from the tryptase area to identify extracellular tryptase (Figure 4D). The total area of extracellular tryptase per biopsy section was higher in EoE than control (Figure 4E, p = 0.0016, n = 9 EoE, 8 control).

For a third method, we hypothesized that tryptase mean fluorescence intensity (MFI) per mast cell would be significantly lower in EoE than control as a result of mast cell degranulation. MFI of individual mast cells as identified by MC-AI were collected and compared across esophageal location and disease state (Table 2). Tryptase MFI was lower in active EoE (p < 0.0001) and EoE remission (p < 0.0001) than control in the epithelium, papillae and LP (Figure 4F). Of note, on the HSS, average papillary mast cell tryptase MFI correlated with eosinophilic inflammation grade (p = 0.002) and stage (p = 0.018), basal zone hyperplasia grade (p = 0.006) and stage (p = 0.0064), and total HSS score stage (p = 0.014); average lamina propria mast cell tryptase MFI correlated with basal zone hyperplasia grade (p = 0.004) and stage (p = 0.0013) (S3B).

**Table 2.**
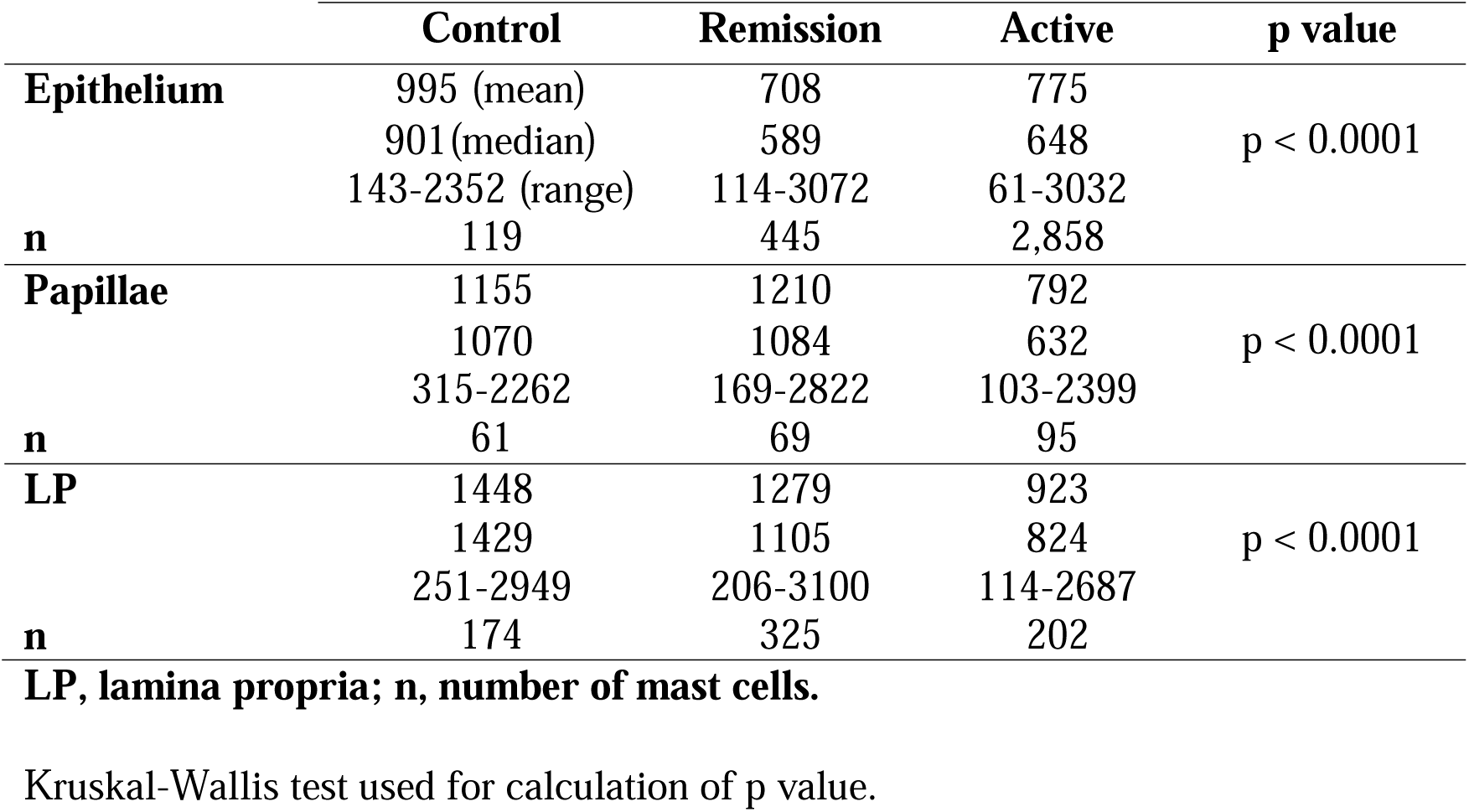
Mast cell tryptase mean fluorescence intensity by location and disease state.

### Correlating mast cell and eosinophil densities in the epithelium and lamina propria

Epithelial mast cell counts positively correlated with peak distal eosinophils per hpf as identified by hematoxylin and eosin (H&E) staining across all disease states (R^2^ = 0.46, p < 0.0001, n = 77 biopsy sections) (Figure S4A) and in active EoE (R^2^ = 0.14, p < 0.03, n = 36 biopsy sections) (Figure S4B). Eosinophil counts were manually enumerated using tissue autofluorescence, which showed a strong positive correlation with mast cell density across all disease states (R^2^ = 0.66, p < 0.0001, n = 39 biopsy sections) (Figure 5A) and in active EoE (R^2^ = 0.49, p < 0.002, n = 17 biopsy sections) (Figure 5B). In contrast, LP mast cell density negatively correlated with eosinophil density across all disease states (R^2^ = 0.26, p = 0.04, n = 16 biopsy sections) (Figure 5C) and in active EoE (R^2^ = 0.56, p = 0.09, n = 6 biopsy sections) (Figure 5D).

**Figure 5.**
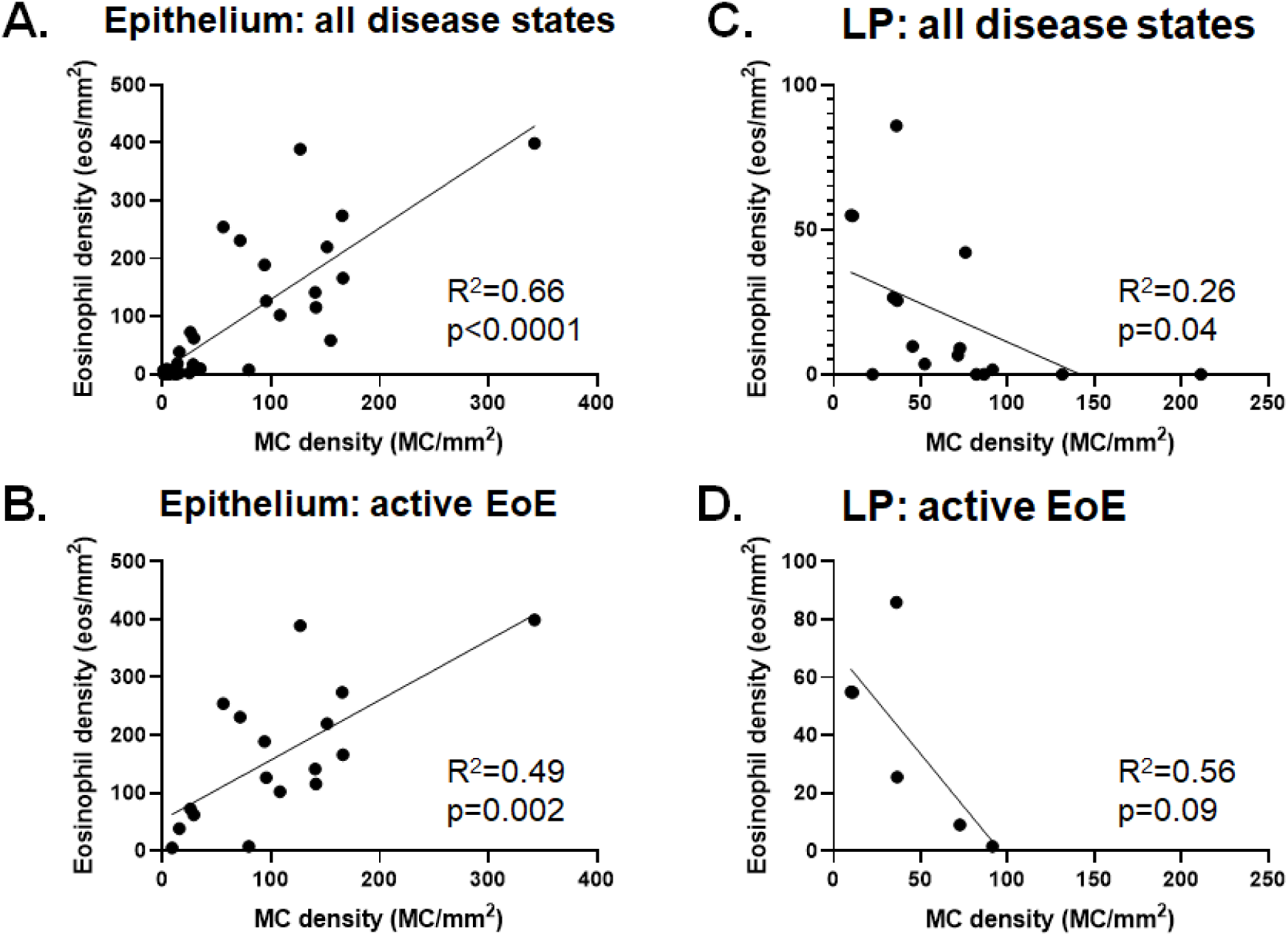
MC-AI–identified MC-eosinophil correlations. Correlation between epithelial eosinophil density (eos, manually enumerated, identified by eosinophil autofluorescence at 532-nm wavelength) and mast cell (MC density) from the same biopsy across all disease states (active EoE, remission EoE, controls) (A) and in active eosinophilic esophagitis (EoE) (Active, B). Correlation between lamina propria (LP) eosinophil density (manually enumerated) and MC density from same biopsy across all disease states (active EoE, remission EoE, controls)(C) and in active EoE (D). Correlations are by simple linear regression.

### Mast cell and eosinophil spatial distribution in the epithelium

Next, we sought to characterize the spatial distribution of mast cells in the esophageal epithelium, as it appeared from biopsy sections that epithelial mast cells were located closer to the basement membrane in control patients and closer to the luminal surface in patients with EoE (active, remission) (Figure 6A). To quantify, we measured the distance of each mast cell from its center to the basement membrane; mast cells were significantly further from the basement membrane in active EoE than in control or EoE remission (Figure 6B).

**Figure 6.**
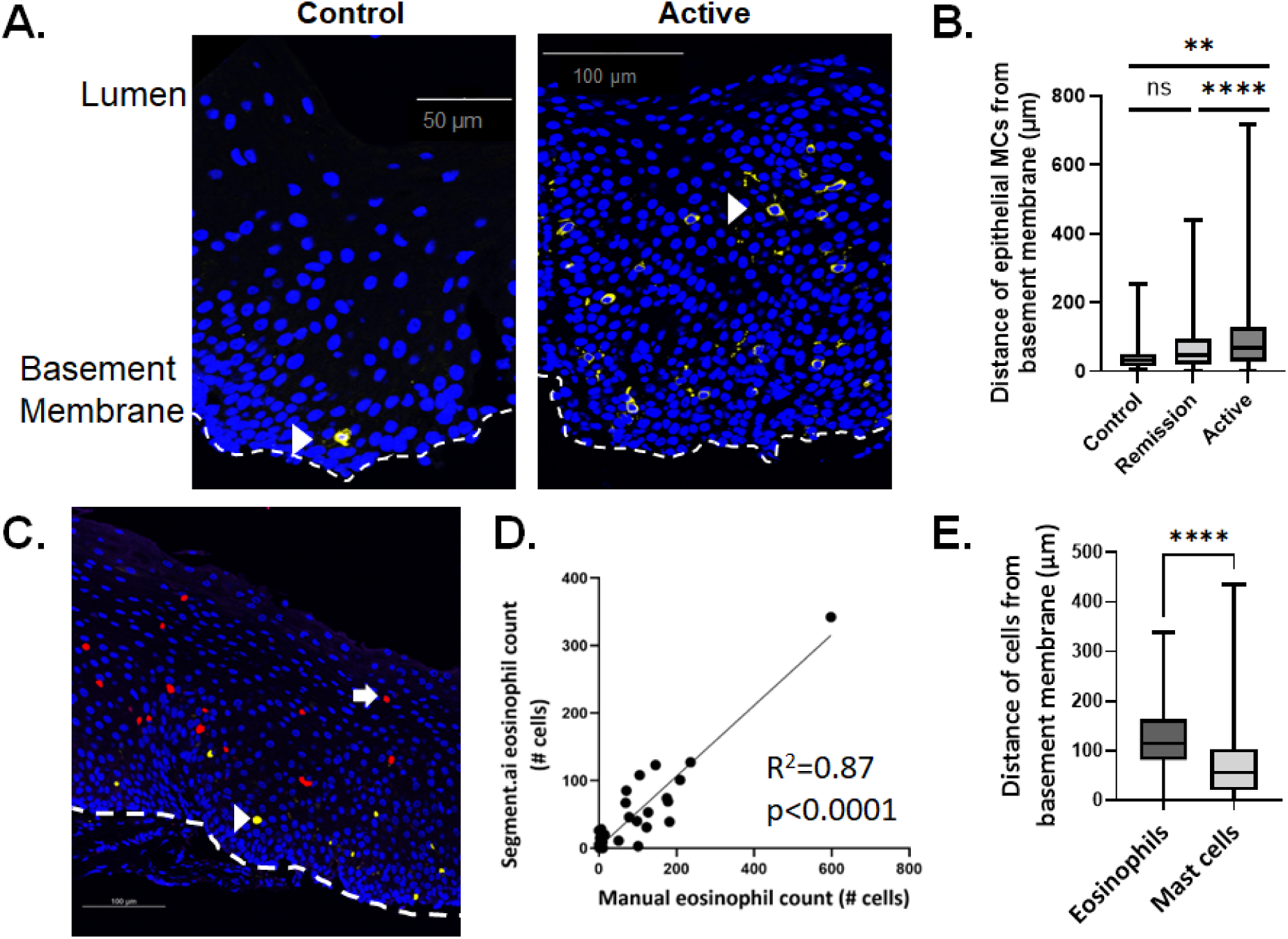
MC and eosinophil epithelial distribution. A. Digitally stitched whole biopsy immunofluorescence of human esophageal tissue in control (left) and active eosinophilic esophagitis (EoE)(Active, right) patient, multiple 40X high-powered fields. Nuclear (blue) and MC-AI–identified mast cell (MC, white) staining. White dashed line: basement membrane. Arrowheads: MCs. B. Distance of individual MCs as identified by MC-AI from the basement membrane in control (n = 20 MCs), EoE remission (Remission, n = 289), and active EoE (n = 514) patients. C. Immunofluorescence of human esophageal tissue in active EoE. Arrow: Eos-AI–identified eosinophils (eos). Arrowhead: MC-AI–identified MCs. D. Correlation between manual and Segment.ai counting of eosinophils (n = 39 biopsy sections) by simple linear regression. E. Distance of eosinophils (n = 438 cells) and MCs (n = 501 cells) from the basement membrane as identified by Eos-AI and MC-AI, respectively. Cell counts were derived from n = 7 patient samples. Boxes represent 25-75% of values; lines represent 1-100%. Statistical difference between the groups *p < 0.0332, **p < 0.0021; ***p < 0.0002, ****p < 0.0001, ns = not significant.

In active EoE, eosinophils appeared to be located further from the basement membrane than were mast cells (Figure 6C). To quantify, we trained Segment.ai to recognize eosinophils (see methods). The Eos-AI protocol–identified eosinophils correlated with manually identified eosinophils (p < 0.0001) (Figure 6D). Eosinophils were found to be further from the basement membrane than were mast cells in active EoE (p < 0.0001) (Figure 6E).

## Discussion

EoE has a strong Th2 signature with evidence of epithelial mastocytosis, but the inability to identify mast cells by H&E staining makes studying these tissue-specific cells difficult. Here, we aimed to further study the spatial distribution and characteristics of esophageal mast cells in EoE by training a machine learning software (Segment.ai) to identify and characterize mast cells. This MC-AI protocol confirmed epithelial mastocytosis, found a decrease in LP mast cells in active EoE, and identified a high density of mast cells in esophageal papillae. The papillae-residing mast cells were shown to dramatically decrease in active disease and negative correlate with epithelial mastocytosis, suggesting that papillary mast cells may be a reservoir for epithelial mast cells. These papillary mast cells were largely CD34 negative, suggesting that they are not mast cell progenitors from the peripheral circulation. Both epithelial and papillary mast cells associated with histologic abnormalities in EoE. Additionally, a second protocol, Degranulated MC-AI, identified mast cell degranulation and determined increased mast cell degranulation in the epithelium, papillae, and LP in active EoE.

The NIS-Elements NIS.ai software contains different features that allow enhanced acquisition, visualization, and analysis^20^. One such feature is Segment.ai, which uses supervised machine learning to identify features of interest. Segment.ai was separately trained to identify mast cells and their degranulation status, and these trained protocols were named MC-AI and Degranulated MC-AI. Combined with high resolution, digitally stitched whole biopsy section IF imaging, these protocols allowed elucidation of novel details regarding mast cell characteristics in human esophageal tissue, including accurately enumerating mast cells and their degree of degranulation. Mast cells are tissue-resident cells that are known to circulate in the blood as mast cell progenitors and terminally differentiate in tissue^21,22^, which have made them difficult to study.

These MC-AI protocols provides a novel way to characterize these tissue-resident cells on a single cell level along with their spatial distribution without relying on visual inspection and manual enumeration. Although MC-AI protocols captured information, there were limitations, such as these protocols likely not capturing every mast cell, some identified cells potentially not being true mast cells, and some captured cells potentially not being whole (MC-AI) or degranulated (Degranulated MC-AI) mast cells.

Esophageal papillae are highly vascular regions of the LP that protrude into the epithelium^17^. We observed that the far-red channel had whole tissue autofluorescence that accurately distinguished the epithelium from LP, which was confirmed by E-cadherin staining. By using this autofluorescence, papillae were manually traced on esophageal biopsies, which helped to delineate mast cells that resided in the papillae versus the epithelium. After separation of the papillae from the epithelium and LP, mast cell density was shown to be highest in the papillae compared to the epithelium and LP in control patients. In EoE (active and remission), this density decreased as epithelial mast cell density increased. Furthermore, a negative correlation was found between papillary mast cell density and epithelial mast cell density, suggesting that papillary mast cells are a possible source of epithelial mast cells. Some of the decrease in density could be due to increase in papillae size, which is documented in EoE, but the fold change in mast cell density is larger than the increase in area (data not shown). Previous work from our laboratory showed that there is *in situ* proliferation of mast cells in the epithelium in EoE^14^. The data presented here show that epithelial mast cells may originate from the mast cells in the esophageal papillae. This population of mast cells may be related to the transitional population of mast cells that have been found through single-cell RNA sequencing analysis in nasal polyps^23^.

Given the vascular nature of papillae, it is conceivable that papillary mast cells are mast cell progenitors that have migrated into local tissue, as mast cells are tissue-resident cells that are known to circulate in the blood as mast cell progenitors and terminally differentiate in tissue^21,22^. However, there were no CD34+ cells found in esophageal tissue other than the endothelium. We detected no CD34+ papillary mast cells, but because the number of mast cells analyzed was small, we can only state that at least 90% of these cells are not progenitors. Mast cell progenitors are known to be CD34+^24^, but there is mixed evidence regarding CD34 expression on mast cells in tissue^25^. One study did not detect CD34 expression on mast cells in nasal polyps by flow cytometry^23^, whereas another identified CD34+tryptase+ mast cells in lung alveolar parenchyma of children with viral lower respiratory tract infections^26^.

MC-AI demonstrated increased mast cell density in active EoE, which confirms previous findings of epithelial mastocytosis^6,8^. Previous data regarding mast cell density in the LP was mixed, with one study showing no change in EoE^8^ and one showing a decrease^27^. Our data demonstrated a decrease in LP mast cell density during active EoE. Further examination of esophageal mast cells showed a more luminal distribution of mast cells from the basement membrane in active disease compared to control.

To investigate the potential effect of mast cells on histologic features of EoE, linear correlations were performed between epithelial, papillary, and LP mast cell density and HSS scores. Epithelial and papillary mast cell density, in contrast to LP mast cell density, correlated with eosinophilic inflammation, basal zone hyperplasia, LP fibrosis, and total HSS score. These findings suggest that papillary mast cells represent a distinct population of mast cells that likely contributes to EoE disease pathology.

Mast cells may be contributing to pathology through degranulation; accordingly, Segment.ai was used to develop Degranulated MC-AI. The density of degranulated mast cells (both Degranulated MC-AI and manually enumerated) was higher in EoE that control, which is consistent with previous data^6^. Our secondary approach of examining degranulated mast cells by enumerating the total amount of tryptase area from the total area of mast cells as identified by MC-AI also demonstrated increased degranulation in EoE compared to control. To further substantiate these findings, we reasoned that the MFI of cell-associated tryptase should be inversely proportional to the degree of MC degranulation. Indeed, tryptase MFI was lower in mast cells in the epithelium in active EoE than control. Interestingly, tryptase MFI was lower in the epithelium in EoE remission as well, suggesting persistent degranulation, consistent with recent single-cell RNA sequencing of esophageal mast cells^14^. Furthermore, the presence of mastocytosis in the epithelium in EoE remission suggests a pathologic role of mast cells even in disease remission. Although the mast cell density in the LP did not increase in EoE, tryptase MFI was significantly lower in EoE remission and even more so in active EoE, suggesting mast cell degranulation in the LP in both active EoE and EoE remission, which may contribute to LP fibrosis.

The characterizations enabled by the MC-AI and Degranulated MC-AI protocols revealed intriguing patterns of mast cell correlation and distribution. Mast cell correlation with basal zone hyperplasia has been previously described^7^. Concurrently, mast cells were found to reside near the basal layer in control esophageal epithelium but were located more luminally in active EoE esophageal epithelium. This localization potentially is related to the basal zone hyperplasia seen in EoE, as well as regional production of CCL26 and other chemokines by the suprabasal epithelium^16^. Mast cell density was found to moderately correlate with peak distal eosinophils/hpf. Segment.ai was trained to identify eosinophils by autofluorescence, and eosinophils were found to be distributed further than mast cells from the basement membrane.

One limitation of this study is the lack of LP in some of the esophageal biopsy sections. As seen in Figure S1A, the amount of LP area is significantly higher in active disease than EoE remission or control biopsy sections.

In conclusion, we have performed an in-depth spatial analysis of esophageal mast cells in EoE by developing a new method that combines high-resolution, whole biopsy section immunofluorescence images and machine learning–based mast cell identification protocol (MC-AI). We identified a new population of papillary mast cells that was present in high density in control, decreased significantly in EoE, negatively correlated with epithelial mast cells, were

CD34 negative, and correlated with features of disease histology. Taken together, these evidence further suggest the pathologic role of mast cells in EoE and have potential implications for anti-mast cell therapy in EoE. Furthermore, the development of MC-AI is an original usage of artificial intelligence which has the potential to advance the understanding of mast cell associated inflammatory diseases, including eosinophilic esophagitis^28–30^.

## Abbreviations

(EoE): eosinophilic esophagitis
(MC): mast cells
(AI): artificial intelligence

## Acknowledgements

We thank the Cincinnati Children’s Confocal Imaging Core and the Digestive Diseases Research Core Center for their assistance and Shawna Hottinger for editorial assistance

## Supplementary Figures

**Figure S1.**
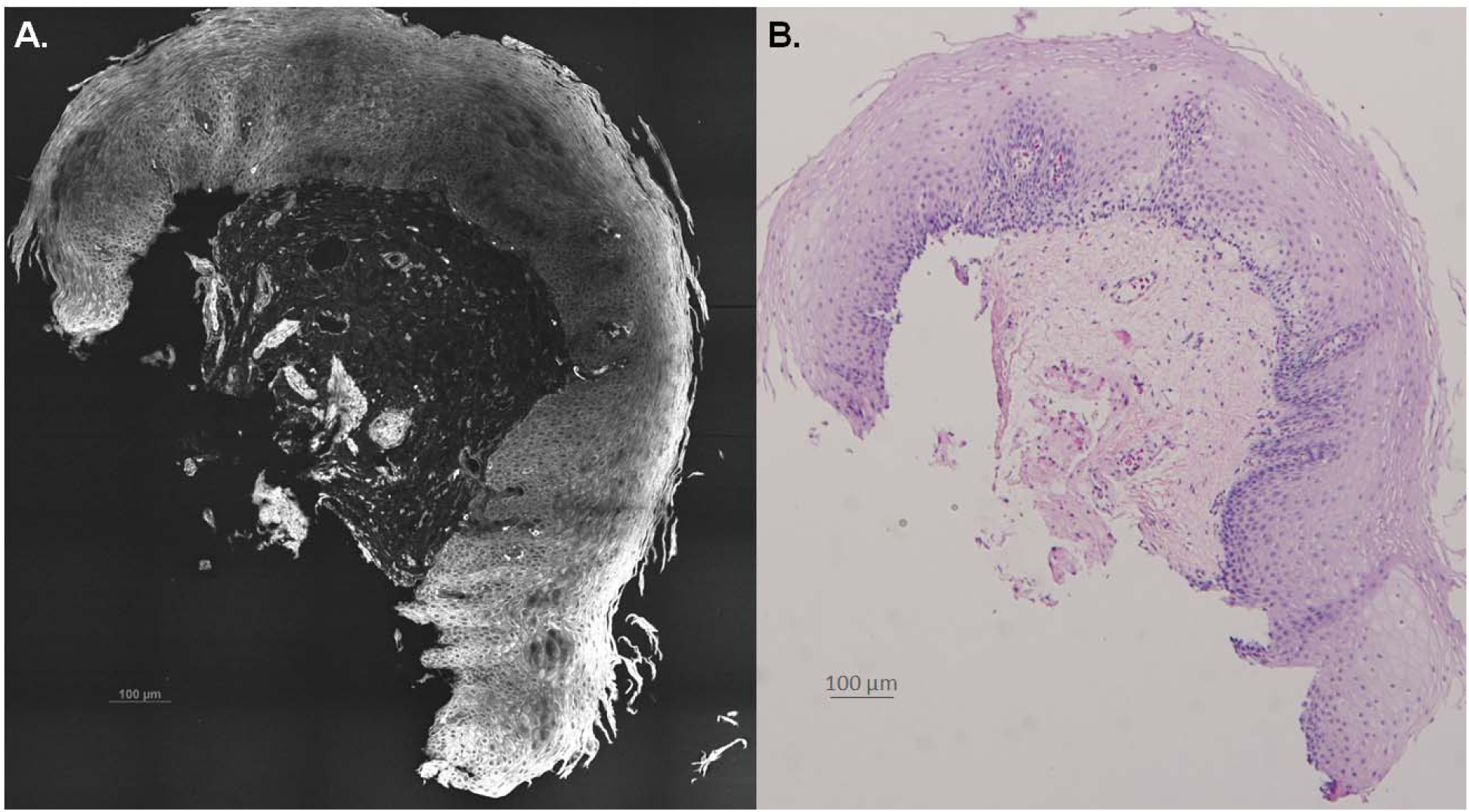
Whole biopsy section. A. Digitally stitched whole biopsy immunofluorescence (IF) of human esophageal tissue, multiple 40X high-powered fields from one representative biopsy section. Tissue autofluorescence (white). B. Histology slide from same biopsy showing tissue architecture.

**Figure S2.**
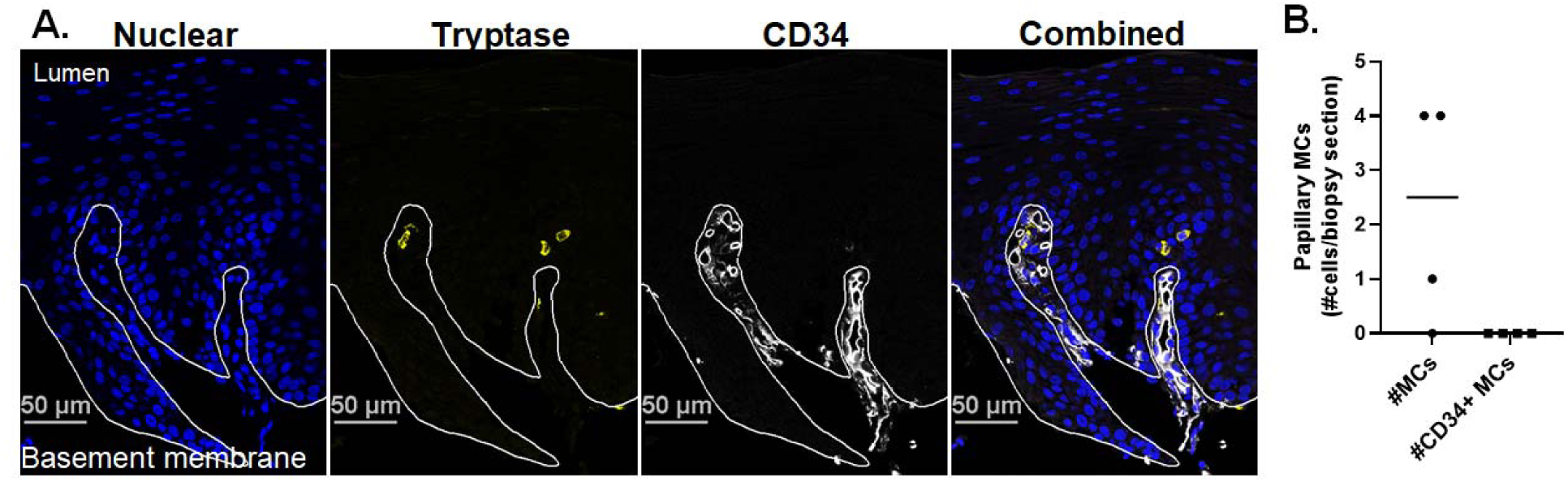
CD34+ MCs in esophageal epithelial tissue in EoE. A. Immunofluorescence of nuclei (left, blue), mast cell (MC) tryptase (middle left, yellow), CD34 (middle right, white), and combined (right). Image is a crop of a digitally stitched image of composite overlapping 40X images from one representative biopsy section. The dashed white line represents the basement membrane. B. Number of epithelial mast cells (MCs) as identified by MC-AI and number of CD34+ epithelial MCs, n = 4 biopsy sections. Markers represent individual biopsy sections; bars represent the median.

**Figure S3.**
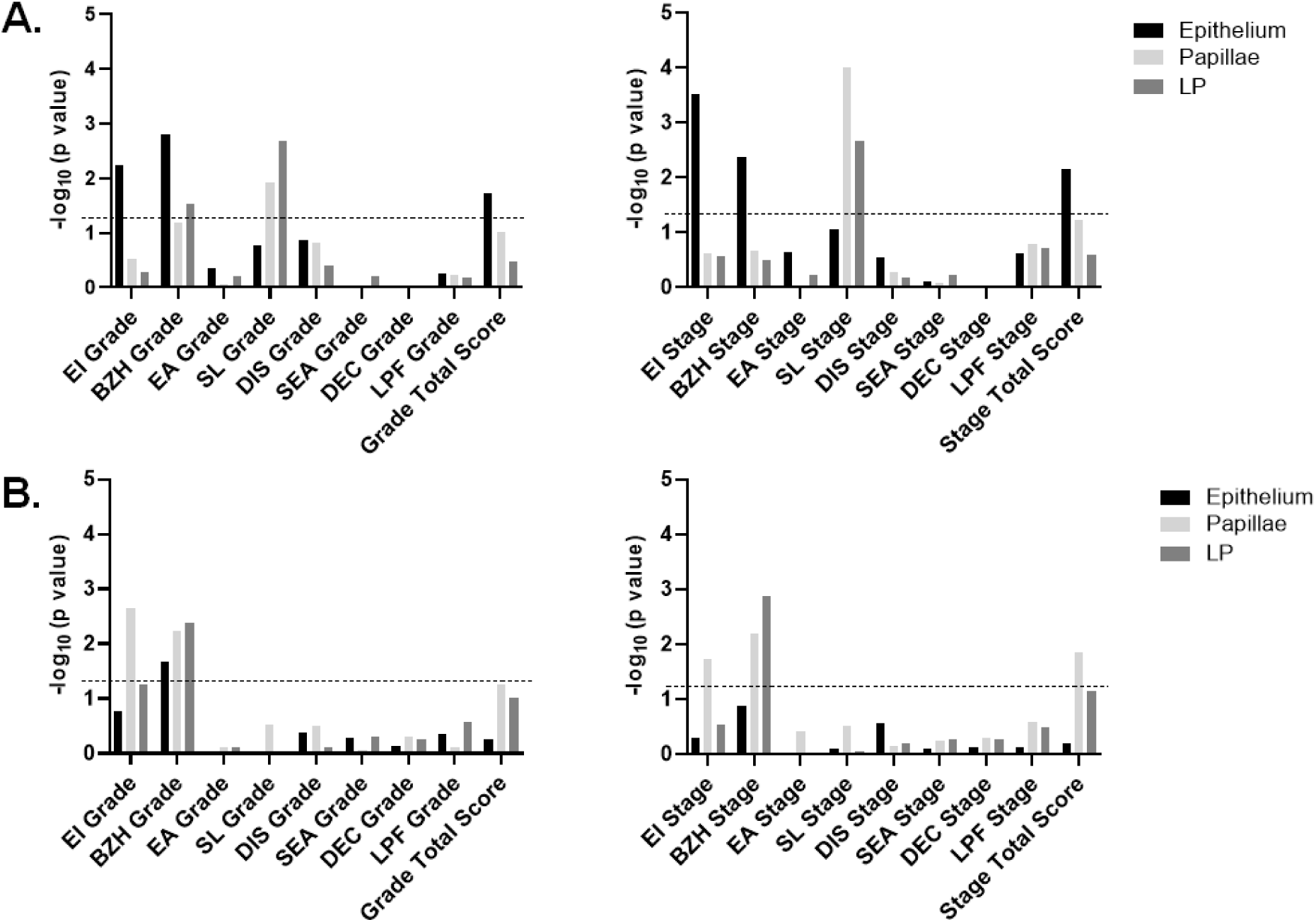
Correlation of Eos-AI–identified eosinophil density and tryptase mean fluorescence intensity (MFI) with Histology Scoring System (HSS) scores across all disease states (-log 10–adjusted p values). A. Correlation of Eos-AI identified eosinophil density with HSS grade scores (left) and stage scores (right). B. Correlation of MC-AI identified MC tryptase MFI with HSS grade scores (left) and stage scores (right). Dotted line represents -log 10– adjusted p value of 0.05. Simple linear regression was used for analysis. LP, lamina propria, EI, eosinophil inflammation, BZH, basal zone hyperplasia, EA, eosinophil abscess, SL, eosinophil surface layering, DIS, dilated intercellular spaces, SEA, surface epithelial alteration, DEC, dyskeratotic epithelial cells, LPF, lamina propria fibrosis.

**Figure S4.**
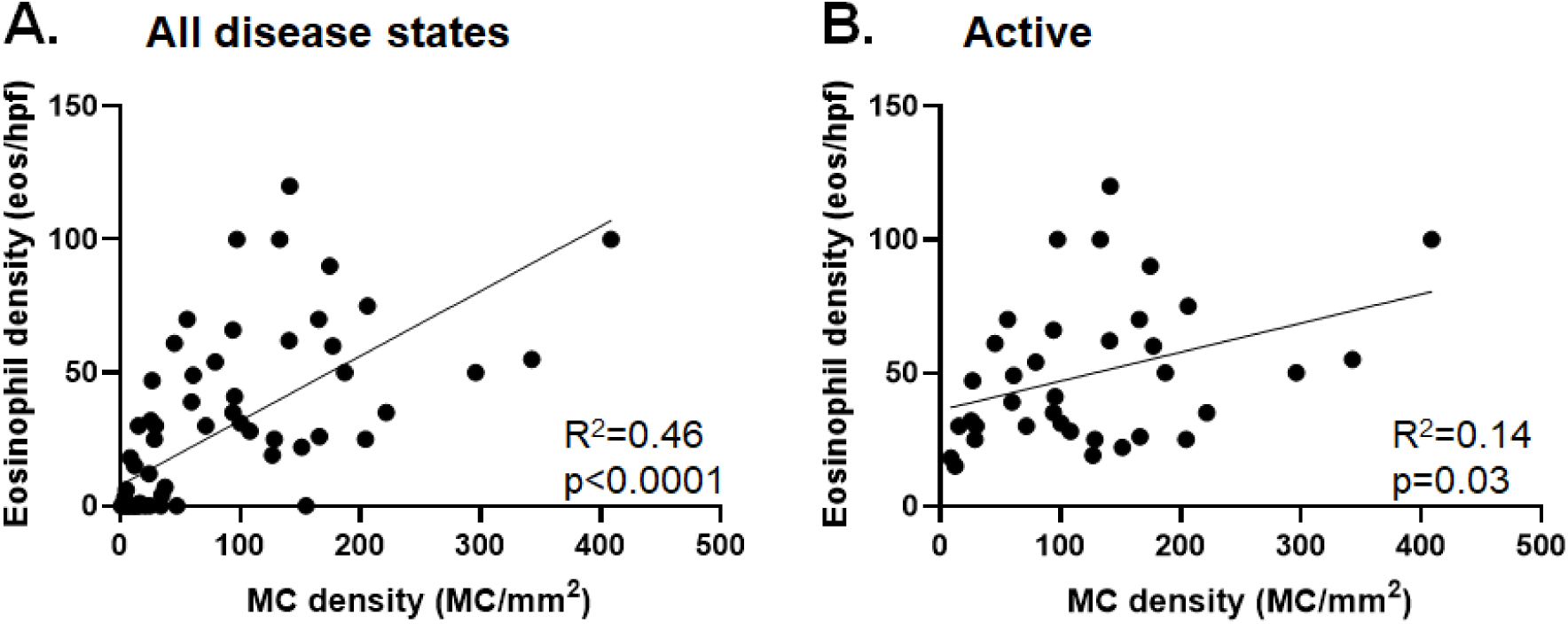
MC-AI–identified MC-eosinophil correlations. Correlation between peak distal esophageal eosinophil counts per high-powered field (hpf) and epithelial mast cell (MC) density across all disease states (A) and in active EoE (Active, B). Correlations are by simple linear regression. Markers represent individual biopsy sections (n = 77 in A and n = 36 in B).

